# Quantifying ethical tradeoffs for vaccine efficacy trials during severe epidemics

**DOI:** 10.1101/193649

**Authors:** Steven E. Bellan, Juliet R.C. Pulliam, Rieke van der Graaf, Spencer J. Fox, Jonathan Dushoff, Lauren Ancel Meyers

## Abstract

**Background:** During emerging epidemics of highly fatal diseases, rapid development and testing of new vaccines may be critical to curbing transmission and saving lives. However, the design of vaccine efficacy trials in such contexts may face considerable logistical, epidemiological, or ethical impediments. Three different vaccine efficacy trials were conducted during the 2014-2016 Ebola virus epidemic in West Africa, each with different designs. At the time, there was vigorous debate on the tradeoff between a trial’s ability to yield information of scientific and societal value versus the perceived ethical dilemma of withholding potentially life-saving vaccines from control participants. Whereas the scientific value of a trial is often estimated in terms of statistical power, speed, and rigor, we lack similar metrics for the ethical costs of withholding interventions.

**Methods and Findings:** Here, we introduce a conceptual framework that fills this gap and allows quantitative assessment of both the scientific value of a study and the risks incurred by trial participants. We show that even untested vaccines against severe diseases may be probabilistically beneficial—i.e. after accounting for realistic uncertainty in their safety and efficacy, trial participants are expected to be better off vaccinated than not. While accounting for this uncertainty, we estimate trial participant risk under a hypothetical, idealized vaccine rollout scenario and compare it to risk under various candidate trial designs, in order to elucidate specific quantitative tradeoffs between cumulative risk to trial participants and information gained. Through an illustrative simulation example, we highlight specific trial-design modifications that allow for conscientious balance between minimizing participant risk and acquiring information of societal value. These include modifications that affect the speed with which a trial would detect an efficacious vaccine (greater sample size or enrollment rate, interim analyses, or risk-prioritized vaccine rollout), which leads to earlier vaccination of control participants should the vaccine be efficacious, and those that systematically limit the risk “spent” by unvaccinated individuals (e.g., providing vaccination to controls after a delay, or presumptive vaccination of subjects above a risk threshold).

**Conclusion:** We advocate this conceptual approach as a means of clarifying the tensions between opposing viewpoints and facilitating transparent discussion to aid ethical and efficient responses to future emerging epidemics.

## Introduction

The West African Ebola epidemic stimulated debate on the comparative ethics of various study designs for Phase III efficacy trials of therapeutics and vaccines. Energetic debate focused on the conflict arising between the combined scientific and societal aims of research, and the welfare of individual trial participants. One camp argued that fast and rigorous identification of safe and efficacious vaccines and therapeutics for epidemic control was the most important criterion. This group supported individually randomized controlled trials (RCT) as the fastest and most statistically efficient study design to evaluate safety and efficacy [1,2]. A second group argued that withholding a promising experimental intervention from control participants at high risk of infection and death may fail to satisfy the principle of clinical equipoise [3]–originally defined as genuine uncertainty in the expert community regarding the *net-preferential* trial arm (i.e., which arm will yield better participant outcomes) [4]. These concerns led to proposals for non-traditional trial designs, including a ring vaccination trial, the use of delayed vaccination instead of control arms, and a stepped-wedge cluster trial (not implemented) [5,6].

Here, we develop a framework that positions these opposing viewpoints within the context of a continuous, quantitative spectrum. In doing so, we highlight design choices that achieve middle ground between them. By clarifying tradeoffs between intervention evaluation and providing benefits to trial participants, we hope to facilitate more efficient and transparent dialogue surrounding trial ethics during future outbreaks of highly fatal pathogens. Here, we focus on assessment of preventive interventions, like vaccines, rather than on therapeutic interventions, which introduce separate logistical and ethical issues.

## Quantifying Expectations based on Prior Evidence and Expert Opinion

It is ethically problematic to withhold a proven life-saving intervention for a condition under study from trial participants, particularly if the intervention is inexpensive and readily accessible [7]. But when might withholding an *unproven* intervention be considered ethically problematic? Several experts argued that it was unethical to withhold experimental Ebola vaccines from trial participants because the anticipated risk of vaccine-related severe adverse effects was extremely unlikely to counterbalance even a conservatively low promise of efficacy [3,8,9], particularly given that interventions would have already emerged with promising safety profiles from Phase I-II trials. Clearly, not all untested interventions are equivalent, and the information available prior to a Phase III clinical trial informs both the perceived value of conducting a trial and expectations with regard to the intervention’s safety and efficacy. A disease’s severity [10] and, for preventative interventions like vaccines, the risk of infection must also be weighed when assessing the expected impact of an experimental intervention. Here, we use the statistical meaning of the word “expectation”–i.e. the expected value (average) of a quantity across uncertainty therein. For an efficacious vaccine, the “net benefit” of vaccination is the decrease in risk associated with vaccination, adjusted for increase in risk associated with adverse events. The argument that withholding vaccines would be unethical rested on the assumption that vaccination would be net-beneficial. For unproven interventions, we refer to the expectation of a net benefit as “probabilistically beneficial” (as opposed to the “demonstrated” benefit from a proven intervention).

It is worth examining this implicitly quantitative argument more closely. Uncertainty in an intervention’s promise can be characterized using probability distributions, which can flexibly describe the range and weight of expert opinion (Figure 1). A Bayesian *prior* is a probability distribution that reflects prior belief (in this case, based on expert opinion) about the probability that an uncertain quantity lies in a given range. For instance, a prior on vaccine efficacy illustrates the expert community’s degree of optimism regarding a candidate vaccine before an efficacy trial, as informed by Phase I-II or pre-clinical studies and other relevant expert experience. The prior probability assigned to the focal parameter (e.g. efficacy) falling in a given range is equivalent to the area under the prior curve within that range (with total area under the curve thus totaling to 1).

**Figure 1.**
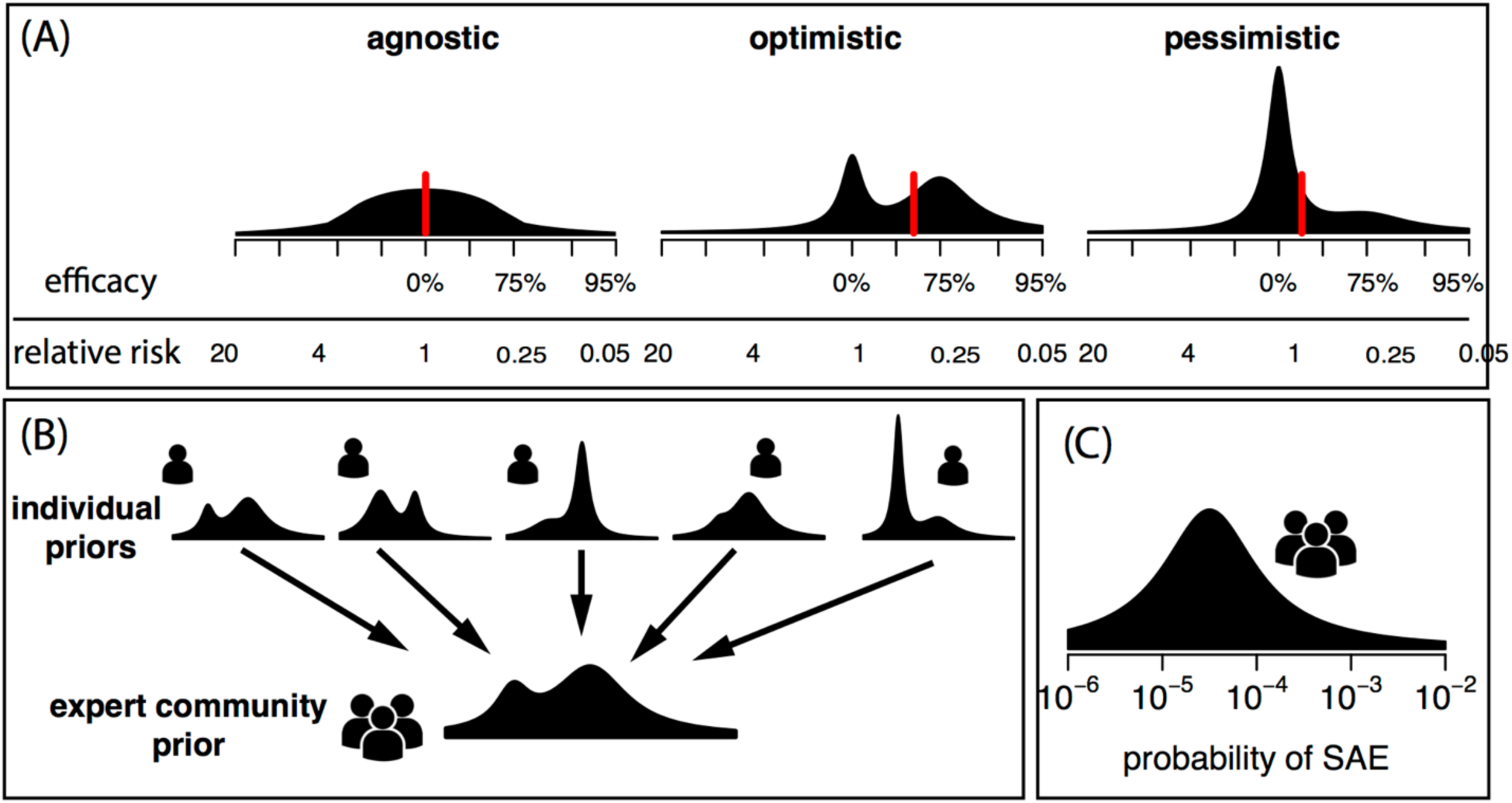
Conceptualization of an intervention’s efficacy and safety using prior probability distributions. Expert opinions on an intervention’s anticipated efficacy prior to a Phase III efficacy trial can be summarized using Bayesian prior belief distributions, which show the probability density an expert would assign to different efficacies (A). Different prior distribution shapes can reflect opinions that are agnostic (equal belief that the intervention may increase or decrease risk), optimistic but conflicted (gives some probability to having a strong efficacy and some to having a small effect in either direction), or pessimistic but conflicted (little probability of a strong protective effect, large probability of negligible effect). Red vertical lines display the mean, or expected values, of these prior distributions. These distributions can, in theory, be collected from multiple individuals and combined to yield prior belief distributions for the whole expert community (B). While this is not easily performed in practice, it is a useful theoretical construct to think about prior opinions and trial ethics. (C) Prior distributions can be similarly created for other aspects of interventions, such as the risk of a serious adverse event (SAE).

Precise quantification of expert opinion into a prior belief distribution will often be challenging, and experts may exhibit bias. Nonetheless, this conceptualization allows for useful thought experiments and methods do exist for eliciting priors from an expert community [11]. With these priors, one can account for realistic uncertainty in intervention safety and efficacy when estimating traditional metrics of a trial’s information value, such as power, speed, and bias. For instance, estimating trial power integrated over an uncertain efficacy yields the probability of a positive finding while accounting for uncertainty in whether the intervention works. In contrast, traditional power estimation conditions on a single assumed efficacy. Under this framing, one can also estimate the effect of withholding an intervention on control participants while accounting for uncertainties in intervention safety and efficacy, as well as in infection risk and case-fatality rates.

## Probabilistic Benefit versus Therapeutic Misconception

The expected efficacy of a vaccine, given uncertainty in expert opinion, can be calculated as the average of the prior distribution on efficacy (Supplementary Text S1). The expected benefit of vaccine protection is a combination of this average efficacy, the expected infection risk, and the expected disease severity. An unproven vaccine can be probabilistically beneficial when infection risk and disease severity are very high, even using conservatively low estimates of the vaccine’s promise (Figure 2 and Figure S1). This is because the expected benefits far outweigh any realistic expectation of risk from the vaccine (see Supplementary Text S1 for further discussion). An unproven intervention that has a 30% chance of 100% efficacy is equally beneficial to a proven intervention that has a 100% chance of 30% efficacy against the same highly fatal disease. Thus, ethical concerns regarding the withholding of probabilistically beneficial interventions may be valid. A less-absolute dichotomization between proven and unproven interventions is useful in many circumstances [12], including when thinking about vaccines against highly fatal, novel diseases.

**Figure 2.**
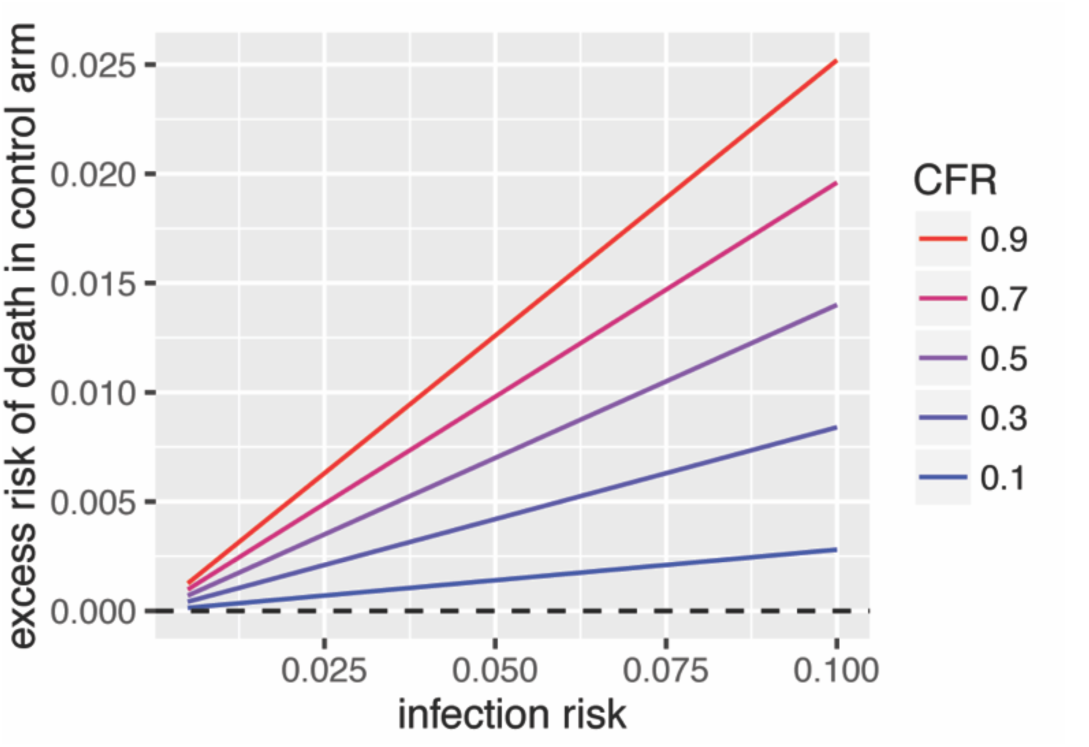
Anticipated excess risk in the control arm. The excess risk of death in the control arm vs a vaccine arm as a function of trial subjects’ infection risk (x axis) and a disease’s case fatality rate (color; CFR), accounting for the expert community’s overall estimate of a vaccine’s efficacy. This example assumes a simple efficacy prior in which experts believe the vaccine has a 50% chance of 70% efficacy and a 50% chance of no efficacy, for an average anticipated efficacy of 28%. The y-axis here can also be interpreted as the magnitude of anticipated adverse side effect risk (on the mortality risk scale) necessary to counterbalance anticipated vaccine efficacy such that there would be no difference in risk between arms. Figure S1 considers wider range of infection risks, serious adverse event risks, and prior assumptions on vaccine safety and efficacy. As noted in the text and Figure 2, we advocate against using uncertainty in the net-preferential arm (the original definition of clinical equipoise) as an ethical goal in itself; however, the expected difference between arms (accounting for uncertainty) may be a good first approximation for the amount of risk spent (Figure 4) by control arm participants (see section on Counterfactual Risk-Benefit Differences).

The notion of probabilistically beneficial interventions has implications for claims that something akin to therapeutic misconception [13] was committed by stakeholders who saw vaccine rollout as a valid goal for Ebola vaccine efficacy trials [9,14]. Subjects under therapeutic misconception (1) mistake the intent of research for that of care, or (2) overestimate the anticipated benefits of an experimental intervention [15]. Rid and Miller suggest that investigators made both these mistakes [13]; however, (2) is not necessarily a misconception if an intervention is probabilistically beneficial, as possible in this context. Further, many agree that (1) has been overstated or misunderstood [16]; trials may reasonably have the dual goals of creating generalizable knowledge and of providing care [17]. Thus, if rollout is probabilistically beneficial, it is reasonable to construe a trial as having the simultaneous goals of intervention evaluation and rollout.

## Withholding interventions: An ethical dilemma?

We can also ask: under what circumstances is withholding a (demonstrably or probabilistically) net-beneficial intervention unethical? For the last several decades, clinical equipoise has served as the principle underlying the argument that withholding benefits from trial participants is unethical. The intent of clinical equipoise is to prevent participant outcomes from being inappropriately subordinated to scientific or societal goals, while still allowing investigators to conduct clinical research with randomized allocation [4]. The idea that there should be uncertainty in the net-preferential arm is related to two other goals related to this uncertainty: (1) the *utility component* requires sufficient uncertainty amongst the expert medical community such that randomization of subjects to two or more trial arms could provide valuable information to society, and (2) the *care component* requires that randomization to these arms not violate the obligation to provide competent care to participants [18]. Competent care means care that lies within the standard of care, is accepted within the expert community, and that does not withhold known superior treatments for the condition under study [19]. But both the nature and extent of investigators’ obligation to provide care to their subjects remain widely debated [18–21]. It is not our goal to resolve that debate here. Rather, we proceed on the assumption that stakeholders are at least obligated to *consider* approaches to maximizing participant outcomes with available resources [22].

The original definition of clinical equipoise does not accurately reflect its application in practice. A simple example illustrates the problem: imagine a proposed trial having arms A and B, with experts in consensus that A is substantially better than B–i.e. the net-preferential arm is *not* uncertain. To make the net-preferential arm more uncertain, investigators could theoretically *worsen* the care in A such that the expected outcomes are more similar to those in B. Clearly, worsening care in one arm benefits no one, and does not make a trial more ethical. Ensuring fairness between arms or, more broadly, uncertainty in the net-preferential arm should not be an ethical goal in its own right because it does not ensure either the utility of a trial or sufficient care to trial participants. Instead, in practice, clinical equipoise may be better characterized as the absence of a conflict between the utility and care components. The outdated focus on uncertainty in the net-preferential arm may explain clinical equipoise’s conspicuous absence from the Declaration of Helsinki [23] and other international guidelines, as well as its rejection as an ethical mandate by many bioethicists and epidemiologists [18].

Another challenge to clinical equipoise as a guiding principle is its binary nature: the two components either conflict or they do not. This dichotomization prevents a more nuanced discussion of trial design modifications that mediate this conflict. We argue that a focus on evaluating tradeoffs between the two components will better facilitate communication among stakeholders. If a proposed study would lead participants to receive care deemed sub-competent by *some* stakeholders, but would provide exceptional information of societal value, these factors need to be evaluated systematically so that a considered decision can be made.

## Counterfactual Risk-Benefit Differences

We propose a new approach to address these challenges: quantifying the conflict between these components and examining how the tradeoff between them can be mediated by specific study-design modifications. Measuring the utility component of a trial can be done using traditional metrics of scientific value–statistical power, trial speed, and bias. The care component, in contrast, has not been measured quantitatively, preventing direct assessment of its tradeoff with trial informativity. We note that these measurements are just a first step; normative judgements are still necessary to balance the societal value of the information acquired and the importance of the care delivered.

To measure the care component, we advocate focusing on *anticipated differences between counterfactual scenarios*, including alternative intervention trial designs, intervention rollout, and inaction (Figure 3), rather than anticipated differences between trial arms. This has several advantages. First, it addresses the question of whether trial stakeholders are doing enough to ensure competent care (as opposed to the less relevant question of whether some people in the trial are getting better care than others). Second, it accounts for key logistical issues that may equally constrain all counterfactual scenarios. Third, it accommodates alternative trial designs that may not include a clear control arm, such as stepped wedge designs, in which all participants receive the intervention but in random order. Finally, in addition to assessments at the population level, it permits consideration of individual-level assessments, which is important because investigators are obligated to provide each participant with competent care [22].

**Figure 3.**
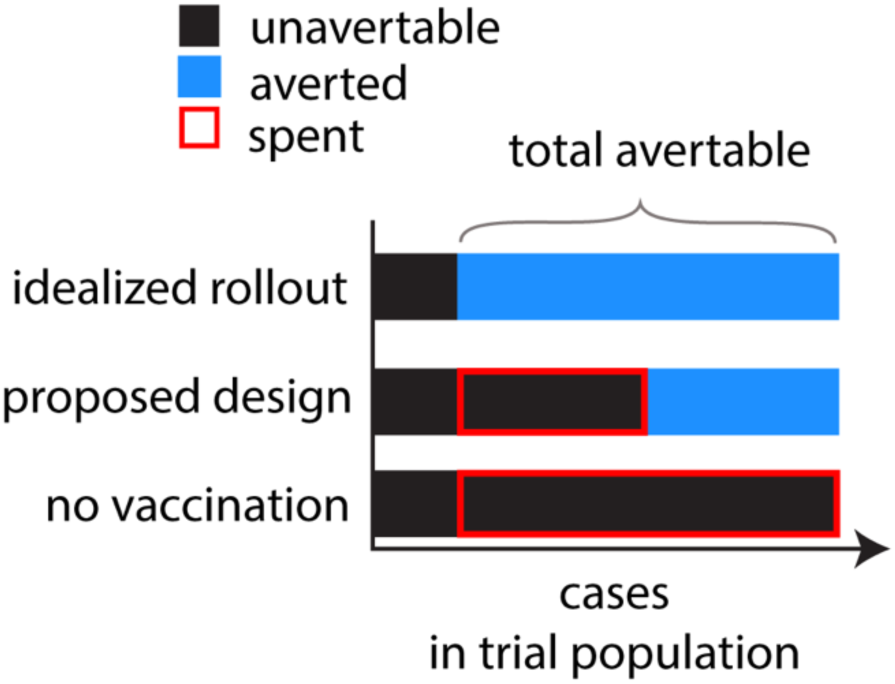
Risk differences between counterfactual scenarios. Participant risk can be partitioned by comparing expected risk (at the trial participant level) or cases (at the trial population level) across scenarios. Avertable risk is defined as the proportion of total risk that could be averted in an idealized intervention rollout in which trial resources are allocated solely towards the goal of averting as much risk as possible (at the trial population level). Averted risk is the subset of avertable risk that would expected to be averted by a specific trial design, while the remainder we denote as risk spent on the information provided by the trial.

The expected amount of participant *risk averted* by a proposed trial can be measured by estimating the anticipated difference between each participant’s expected outcomes in that trial and in a scenario of inaction (i.e., no vaccination). The amount of participant *risk spent* (i.e. avertable risk that is not averted in order to acquire information) can be measured by estimating the anticipated difference between each participant’s outcomes in a proposed trial and an idealized rollout scenario in which investigators allocate their available resources solely towards optimizing participants’ outcomes (for the condition under study), given logistical constraints. This idealized rollout scenario is a useful reference point from which to examine alternative designs; it is not necessarily a goal in itself. Notably, risk spent is not “harm” because it is not additional risk created by the trial (compared to the baseline of inaction), but only additional risk compared to this idealized counterfactual [7]. Finally, while we dispute the notion that uncertainty in the net-preferential arm should be agoal in itself, the expected difference between arms (Figure 2) can still be a useful first approximation of risk spent; this is true when outcomes in the net-preferential arm closely resemble the outcomes of the entire trial population in the idealized scenario. In many cases, however, logistical constraints or other complexities may invalidate this assumption, e.g., because not everyone can be vaccinated simultaneously. In such cases,detailed simulations that account for logistical constraints will provide a better lens into the appropriate counterfactuals.

These counterfactual differences can be contextualized as follows (Figure 3). We call a participant’s risk in the scenario of inaction *total risk*. The anticipated difference between inaction and the idealized scenario is *avertable risk*. Not all risk is avertable because of uncertainty, imperfect vaccine efficacy, serious adverse effects, or logistical constraints on rollout speed. Avertable risk is further divided into that which is in fact averted and that which is spent. Comparing averted to avertable risk prevents overestimating the effect of withholding interventions on trial participants. For instance, an individual’s risk when randomized to control in an RCT should be compared to that individual’s vaccination timing under the idealized rollout scenario rather than to instant vaccination, which might not occur in any realistic scenario (e.g. if that individual lives in an area that would not be prioritized for early vaccination).

Estimating these counterfactual risk differences can be done in four computational steps: (1) fit a sufficiently detailed simulation model to available case data to project realistic epidemic risk trajectories with uncertainty; (2) draw thousands of random vaccine efficacies from a prior distribution that reflects expert uncertainty; (3) for each random efficacy, superimpose a candidate trial design, inaction, and the idealized rollout scenario, in turn, on a simulated trial population; and (4) for each scenario compute trial-level metrics and the average outcome for each participant across these thousands of simulations, taking the differences defined above to calculate risk spent and risk averted. We consider both individual-level metrics (*risk spent* or *averted*) as well as metrics that sum risk across the trial population (*cases spent* or *averted*).

## Design Modifications to Limit Risk Spent: An Ebola vaccine trial case study

Here, we develop an illustrative example based on an Ebola vaccine trial scenario, which demonstrates how design modifications affect risk spent, power, and trial speed. We agree with the consensus that, for the sake of rigor and efficiency, trial design should include individual randomization, double blinding, and placebo control whenever possible [24]. Accordingly, most of the scenarios we consider incorporate these aspects. Our example extends a trial simulation framework previously used to evaluate Ebola vaccine trial plans in Sierra Leone [25]. All scenarios assume a trial population of 10 sites of 200 individuals and assess designs that use either individual- or cluster- (i.e. site-) level randomization. We assume a logistical constraint on speed of vaccine rollout across all scenarios, likely representative of most epidemic scenarios in which immediate vaccination is not feasible. Vaccine efficacy is randomly sampled from a prior distribution intended to represent expert opinion. Similarly, we do not assume perfect knowledge of cluster- or individual-level heterogeneity in risk, or temporal variation therein. We assume cluster heterogeneity is partially predictable based on recent case trends. We also assume that, within an area, role-related heterogeneity in infection risk (i.e. healthcare workers for Ebola) may often be partially understood *a priori*. Further detail is available in Figure S2 and Text S1.

We illustrate the effects of specific design choices via simulation of the following designs: (1) a standard randomized controlled trial (RCT), in which clusters are enrolled in a random order and randomization to trial arm occurs at the individual level within clusters; (2) RCT with interim analyses (int.) that allow for trials to end earlier, (3) RCT int. with risk-prioritized order of vaccine rollout (r.p.) such that high-risk clusters are enrolled first, (4) RCT int. r.p. that presumptively vaccinates and excludes from analysis the 15% highest risk individuals (e.h.r.), (5) RCT int. r.p. using a 60-day delayed vaccine comparator instead of a control arm (d.v.c.), and (6) a stepped wedge cluster trial (SWCT). We examine these designs in both waning and peak epidemic contexts (represented as trial start dates of November 1 and December 1, 2014, respectively), to represent situations when acquiring enough cases for adequate power is and is not a major concern.

The first category of design modifications that limit risk spent are those that increase the speed with which a trial will detect an efficacious vaccine. Because trials roll out interventions to all participants once the intervention has been shown successful, modifications that increase speed *indirectly* reduce the time spent unvaccinated by control participants, reducing their risk spent. Approaches to increase speed include increasing trial sample size or enrollment rates; the use of group sequential designs in which predefined interim analyses allow trials to end earlier; and risk-prioritized enrollment such that the highest risk (and thus most informative) individuals are enrolled earliest. These modifications have distinct collateral effects. Greater sample size or faster enrollment entails greater cost and may be logistically infeasible. As shown in Figure 4, implementation of interim analyses can dramatically increase trial speed, with a consequent reduction in total risk spent when overall risk to trial participants is high (e.g., near the epidemic peak), though this comes at a slight cost in statistical power. In contrast, risk-prioritization is a win-win, resulting in further substantial reductions in risk spent and, depending on the epidemic context, also increasing power [25,26].

**Figure 4.**
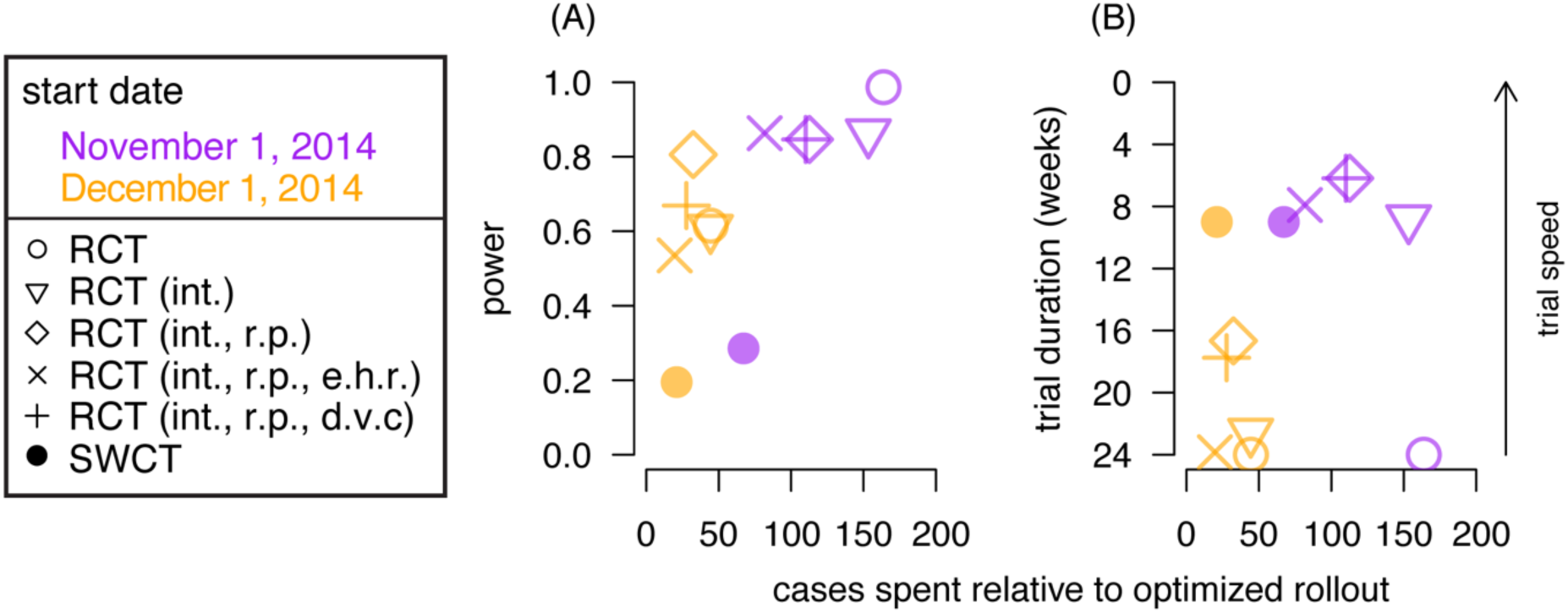
A simulation case study illustrating the effect of trial design and epidemic phase on risk spent for an Ebola vaccine trial in Sierra Leone. Risk spent is characterized here as the average excess number of cases occurring in the trial population relative to the idealized rollout scenario, cumulative over 48 weeks from trial start (“cases spent”). (A) and (B) display the statistical power and mean trial duration, respectively, versus cases spent for various designs and two trial start dates. Simulations use random vaccine efficacy parameters sampled from a prior distribution to account for uncertain vaccine efficacy (0.3 probability of no efficacy, 0.7 probability of uniformly randomly distributed efficacy from 0.5 to 0.9). All individually-randomized trials have a maximum duration of 24 weeks with, if applicable, pre-planned interim analyses based on O’Brien Fleming-like alpha spending rules. All designs are logistically constrained by rollout of vaccine to one cluster per week during randomization and post-trial rollout. SWCT designs end when the last of the ten clusters is vaccinated (after 9 weeks) since there are no unvaccinated individuals left in the trial to which to compare vaccinated individuals. Randomized controlled trials (RCT) may have interim analyses (int.), risk-prioritized order of vaccine rollout (r.p.) to clusters, or exclude the 15% highest risk individuals (e.h.r); they may also use a 60-day delayed vaccine comparator instead of a control arm (d.v.c.). Power is defined as the probability that a trial is successful at detecting statistically significant vaccine efficacy amongst simulations in which efficacy is positive. More detail on trial simulation is provided in Online Supplementary Text S1 and [25].

The second category of design modifications to limit risk spent are those that *directly* limit unvaccinated risk or time. Presumptive vaccination of individuals known to be high risk (e.g. based on their job or contacts) will substantially reduce their risk spent (Figure S4). Of course, those excluded from randomization must be excluded from analysis to prevent bias, resulting in a reduction in power (Figure 4A). The exclusion of high-risk individuals can also substantially increase trial duration, as demonstrated in Figure 4B; in the waning epidemic scenario simulated here, this modification negates the gains in speed that result from incorporation of interim analyses and cluster-level risk prioritization. Importantly, while exclusion of high-risk individuals reduces overall risk spent, the consequential reduced speed of the trial increases the risk spent by lower-risk participants (Figure S5). High risk individuals could alternatively be given the choice of presumptive vaccination or randomization, allowing them to choose whether to spend their risk on information of societal value.

Another direct approach consists of using a delayed vaccine comparator arm (versus an unvaccinated control arm) to systematically limit the time individuals spend unvaccinated [5]. In our case study, this approach has a small effect when transmission intensity is high (i.e. in the peak epidemic context); because a trial is likely to end early if the vaccine is efficacious (with control individuals getting vaccinated), the further reduction in risk spent that results from having a delayed vaccine comparator is negligible. In the waning epidemic scenario, when cases are relatively rare, using a delayed vaccine comparator reduces risk spent but at a small cost to trial speed and power. Delayed-vaccine comparators may be a particularly reasonable method of reducing risk spent when slow trial speed is driven, not by low infection risk, but by long delays between enrollment of each group of individuals or by long delays between infection and the primary outcome, such that participants enrolled early do not continue to spend risk while additional clusters are enrolled [5] or while a lengthy disease natural history elapses (e.g. a vaccine efficacy trial in mothers in which congenital Zika syndrome is the primary outcome), respectively.

The final trial design we consider (SWCT) was proposed for use during the Ebola epidemic because the absence of a control arm was considered an ethical advantage. A SWCT differs from delayed-vaccine designs because in the former, delays in vaccination are driven by logistical constraints, whereas in the latter, they are driven solely by study design. Because the SWCT vaccinates individuals as quickly as possible, subject only to logistical constraints, it appealed to those who argued in favor of vaccine rollout; however, our simulation-based framework highlights that this approach does not minimize risk spent. In order to have scientific value, a SWCT must randomize the order in which clusters are vaccinated. This implies that higher-risk clusters may be randomized to receive vaccination after lower-risk clusters, which differs from both an idealized rollout scenario and a risk-prioritized RCT. In our simulation results, the SWCT appears comparable to the modified RCT study designs in terms of risk spent; however, this result is based on the assumption that the lack of a control (or comparator) arm will allow vaccination to proceed twice as quickly in a SWCT than an RCT (that is, the imposed constraint is the speed with which a new cluster can be reached by the vaccination team). If instead the *individual-level* vaccination rate were similarly constrained across trial designs, the risk spent in a SWCT would be similar to that of an RCT without risk-prioritized vaccination rollout but, as shown previously, the SWCT has far less power than an individually randomized design because of the inefficiency of cluster randomization when risk is spatiotemporally variable [25].

## Discussion

During the West African Ebola epidemic, some argued that it was ethically problematic to withhold experimental interventions from control-arm subjects. Others argued that experimental interventions cannot be presumed efficacious *a priori* and must be rigorously tested before widespread rollout. We provide a quantitative framework that unifies these two seemingly disparate perspectives about vaccine evaluation in the face of a deadly epidemic. Our framework links each argument to a different component of clinical equipoise and provides a pragmatic framework for balancing these perspectives when designing trials. This framework recognizes that, in some cases, conceiving of efficacy trials as having the dual goals of intervention evaluation and deployment is not equivalent to therapeutic misconception. When risk is high, an intervention can have large probabilistic benefits *a priori*, even under conservative assumptions about its promise.

When clinical research includes the dual goals of evaluation and rollout, balancing these goals can be aided by using quantitative metrics. We use traditional metrics of evaluation (power, speed, rigor) and develop a counterfactual framework in which to examine participant outcomes. We advocate appropriately detailed simulation models to help understand how certain design modifications can pragmatically balance between these goals in a specific epidemic and logistical context.

The framework presented here is not intended to replace other considerations regarding informed consent, community engagement, logistics, or resource availability. Rather, this framework aims to complement those considerations in a way that allows various stakeholders to explicitly discuss how they would like to balance the risk spent by trial participants with trial outcomes that may provide societal value for others. Our approach is not prescriptive. We do not recommend a specific balance between these views as the single most ethical. Rather, we argue that discussion of these issues can be rendered more transparent by quantifying the relevant tradeoffs. Simulation studies that aim to quantify the population impact of the rollout of an intervention, if it emerges successfully from an efficacy trial, may be helpful in evaluating the *societal* value of a trial (i.e. beyond measures of its *scientific* value).

Our case study illustrates how this framework could have added clarity to study design decisions during the West African Ebola epidemic. For example, we show that if a trial had been conducted during the peak of the epidemic in the trial population considered, it is possible that high-risk individuals could have been excluded from randomization and vaccinated without appreciable effect on trial speed or power. We show that delayed-vaccine comparators are valuable for reducing risk spent, but only when delayed vaccination is likely to occur substantially before demonstration of efficacy. Finally, we also show that the SWCT does not exhibit its anticipated ethical advantages (i.e., does not reduce risk spent)–unless it is assumed that this design is capable of vaccinating individuals substantially faster than individually randomized designs (because entire clusters are vaccinated versus half-clusters).

This case study does not comprise a comprehensive investigation of all possible study designs for evaluating a vaccine during an emerging epidemic; many other designs are possible and may be preferable for certain settings. Our goal is to provide a unified and pragmatic approach to compare designs within a given clinical and epidemiological context. Some designs present straightforward extensions to those presented here [5,26]. Others may require more challenging comparisons. For instance, comparing designs with different inclusion criteria (and therefore disjoint trial populations) will require counterfactual comparisons that include risk spent by individuals who are in one trial but not in another, complicating the individual-level considerations of competent care. Similarly, the simulations underlying the illustrative examples presented here rely on a relatively simple model of infectious disease risk. The conceptual risk-spending framework, however, can be adapted to transmission models and trial designs of any complexity.

Simulation presumes some level of predictability of infection risk. While accurate and precise prediction of epidemic trajectories is challenging, projections do not have to be perfect to be useful. Further, stakeholders can account for uncertainty in projections by considering several scenarios, or by directly including this uncertainty in trial simulations. Finally, while we advocate simulation as a critical step in planning any trial, a full-scale simulation study of how risk spent trades off with other trial characteristics will not always be necessary. We believe that even framing the questions of how to balance competing trial goals within the proposed framework will help sharpen discussion surrounding trial plans.

While this framework can also be applied to therapeutic intervention trials, there are key differences. First, efficacy and safety priors are likely to be more optimistic for vaccines than for drugs, given their respective track records [27]. Second, the above design modifications are less relevant for therapeutics. Therapeutic effectiveness is generally very sensitive to timing of administration relative to the time of infection or symptom onset, particularly for acutely fatal diseases. Thus, control participants are unlikely to receive benefits from post-trial intervention rollout, such that trial speed will not impact their risk spent. Lack of flexibility in treatment timing also negates the ability of delayed-comparator designs to reduce risk spent. Finally, exclusion and presumptive vaccination of high-infection-risk individuals reduces the power of vaccine efficacy trials but not necessarily their generalizability because vaccine efficacy is unlikely to vary with exposure risk, which is mostly behaviorally driven. In contrast, therapeutic efficacy may vary with disease severity, so evaluating efficacy amongst participants with severe disease may be necessary.

## Conclusion

When trial subjects are at substantial risk of highly fatal infections, experimental vaccines may be probabilistically beneficial, meaning that subjects may be better off receiving an unproven vaccine than not. In such situations, it is rational to conceive of trials as simultaneously having the dual goals of vaccine evaluation and of rollout to protect trial subjects. The conceptual framework proposed here can be adapted to context-specific simulations to examine the tradeoffs between these goals along multiple dimensions of trial design. In doing so, the false dichotomy of whether or not to randomize can be replaced with a higher, more transparent level of discourse that centers on explicitly finding a consensus balance between trial subject health and the societal value yielded by a trial.

## Acknowledgements

We acknowledge Christopher Whalen for useful feedback. We also acknowledge the Texas Advanced Computing Center (TACC) at The University of Texas at Austin for providing high performance computational resources that have contributed to the research results reported within this paper (<u>http://www.tacc.utexas.edu</u>).

## Conflicts of Interest

The authors have declared that no competing interests exist.

## Author Contributions

SEB conceived the project idea, performed the analyses, reviewed the literature, and wrote the first draft. All authors contributed to the development of the project idea, interpretation and presentation of results, and the writing and approval of the final manuscript.

## Funding

SEB was supported by National Institute of Health (NIH) National Institute of Allergy and Infectious Diseases grant K01AI125830. SEB and JRCP were supported by the International Clinics on Infectious Disease Dynamics and Data (ICI3D) program, which was supported by NIH National Institute of General Medical Sciences (NIGMS) grant R25GM102149. SEB, SF, and LAM were supported by NIH NIGMS MIDAS grant U01GM087719. JRCP was supported by the National Science Foundation Rapid Response Research Program (RAPID grant 1515734), the Research and Policy on Infectious Disease Dynamics (RAPIDD) Program of the Fogarty International Center, National Institutes of Health and Science and Technology Directorate, Department of Homeland Security, and the DST-NRF Centre of Excellence in Epidemiological Modelling and Analysis (SACEMA), hosted by Stellenbosch University. JD is supported by the Canadian Institutes of Health Research (CIHR) and the Natural Sciences and Engineering Research Council of Canada (NSERC). The funders had no role in study design, data collection and analysis, decision to publish, or preparation of the manuscript. SEB had full access to all the data in the study and final responsibility for the decision to submit for publication. The content is solely the responsibility of the authors and does not necessarily represent the official views of the funding agencies.

